# Radiation damage behavior of soft matter in ultrafast cryo-electron microscopy (cryo-UEM)

**DOI:** 10.1101/2024.12.20.629524

**Authors:** Yimin Zhao, Chen Qi, Chunhui Zhu, Yun Zhu, Yongzhao Zhang, Tongnian Gu, Huanfang Tian, Wentao Wang, Siyuan Huang, Huaixin Yang, Jianqi Li, Fei Sun

## Abstract

Whether time-modulated pulsed-electron imaging in ultrafast electron microscopy (UEM) can mitigate the electron radiation damage that occurs to samples, is still controversial. The effectiveness of such mitigation effect and relevant potential application in cryo-EM remain to be explored. Herein, we built an ultrafast cryo-EM (cryo-UEM) device based on an ultrafast laser system. Using such equipment and the saturated aliphatic hydrocarbon compounds (C_44_H_90_), the fading curves of diffraction intensity and corresponding critical electron doses (*N*_*e*_) of the samples were carefully measured under different imaging modes, temperatures, imaging dose rates and pulsed repetition rates. Our experimental results show that, the fading curves and *N*_*e*_ values of the C_44_H_90_ crystals are uncorrelated with the imaging electron dose rates and do not show dependence on the dose-rate effect. As the temperature decreased, the *N*_*e*_ values of the sample increased, indicating the cryoprotective effect on radiation damage to the samples. Surprisingly, at a constant temperature, the fading curves and *N*_*e*_ values of the sample in multi-electrons-packet and near-single-electron-packet pulsed modes are all approximately the same as those in conventional continuous electron-beam mode, even when the results are obtained at different pulsed repetition rates. These results show that the time-modulated pulsed electron beam does not seem to mitigate the electron radiation damage that occurs on samples. Our findings offer new insights and experimental basis for the radiation damage behavior of samples under electron beams, and provide guidance and inspiration for elucidating the fundamental principles of radiation damage.

## Introduction

Cryo-electron microscopy (Cryo-EM) has revolutionized the research progress in the field of life science through the “resolution revolution” (1-3), using methods such as sample cryoprotection (4, 5) and low-electron-dose imaging (6, 7) to mitigate the electron radiation damage during the imaging process. However, electron radiation damage is still the main factor limiting the further improvement of cryo-EM resolution, which is usually caused by electron-beam-induced atomic displacement and electronic excitation, and progressively triggers a series of cascading reactions leading to electrostatic charging, heating, radiolysis and diffusion events through inelastic electron-electron or electron-phonon scattering (8, 9). The factors and mechanisms associated with electron radiation damage are usually numerous and complex, with radiolysis originating from long-lived electronic excitation being the primary radiation damage mechanism for both organic and biological samples (10-12). Cumulative ionization damage and bond breaking limit the structural information that can be obtained before the sample destroyed (13). The ultimate bottleneck in cryo-EM technology is inevitably the physical nature of radiation damage to biomolecules during imaging, which is unavoidable by cryo-EM technology and limits the ability to achieve atomic resolution (14-16).

Despite relevant experiment has been reported recently (17), whether ultrafast pulsed electron imaging can mitigate radiation damage in soft matter remains controversial (12, 18, 19). Experiments by Flannigan *et al*. showed that the radiation damage of C_36_H_74_ crystals in ultrafast single-electron-packet pulsed mode was reduced by nearly 2 times compared to continuous electron-beam mode at the same dose rates (17). During radiation damage by cumulative ionization damage and bond breaking, the excitation of each cascade reaction has their own characteristic temporal window (20-22). Flannigan *et al*. believed that the segmented delivery of electrons using time-modulated pulses would allow sufficient relaxation time and sample recovery time for the cascade reaction and subsequent phonon excitation (23, 24), intercepting the occurrence of damage events and decreasing adverse synergistic or composite effects, thus reducing electron radiation damage to the sample to a certain extent. It should be noted that in their study (17), the cumulative electron doses acting on the sample were very low (<0.1 e^−^/Å^2^), and the critical electron doses (*N*_*e*_) of the sample were not obtained, which may be due to the long data collection time of pulsed electron imaging with the ultrafast laser-pulsed system. Glaeser *et al*. suggested that, the claim that radiation damage could be reduced by using pulsed sources rather than continuous sources to deliver the same number of electrons in the same total exposure time was incorrect (25). They thought, to a first approximation, the average time between electrons could be arranged to be the same whether the electrons were emitted from the continuous sources or pulsed sources. As a result, simply reducing the intensity of a standard continuous electron source, rather than using an ultrafast pulsed source, should be much easier to provide a suitably long pause between individual inelastic scattering events for postulated thermal relaxation and “repair”, implying that dose-rate effect should correlate with electron radiation damage to the sample. However, this was not the case in the experimental results of Glaeser *et al*. (26, 27), which showed that dose-rate effect was independent of electron radiation damage.

Whether the dose rate correlates with radiation damage has also been debated for decades, and experimental results are controversial in both cryo-EM and X-ray crystallography (18). Experiments supporting the view that dose rate is independent of radiation damage found that, at the same total doses, lower dose rates were not conducive to high-resolution imaging (26-31), where the fading of diffraction spots was only related to the total cumulative electron doses. Experiments supporting the opposite view suggested that there was dose-rate-dependent damage to the sample and that the critical dose was positively correlated with the dose rates (32-34). Since the mechanism of radiation damage to samples is still not clear (35), the correlation between radiation damage and dose rate remains to be explored.

With the progress of related technologies and methods, ultrafast electron microscopy (UEM) provides new opportunities to address the problem of conformational changes in biological samples (36-40). The combination of UEM with cryo-EM not only enables ultrafast dynamic structural characterization of biomolecules on multiple time scales (39, 41-43), but also promises to be developed as an effective means to explore the electron radiation damage behavior of low-temperature samples through ultrafast pulsed electron imaging modulated by time and electron number. On this basis, by modifying the hardware and software of conventional cryo-EM and introducing ultrafast laser system, we successfully built a biological ultrafast cryo-EM (cryo-UEM) equipment (*SI Appendix*, Fig. S1), and carried out a systematic study on whether the electron radiation damage of soft matter has time-modulated mitigation effect and dose-rate dependence.

Here, we use cryo-UEM to study the radiation damage behavior of C_44_H_90_ crystals at different temperatures and modes. By overcoming problems such as electron-dose fading and collecting experimental data for a long time, continuously and repeatedly, we obtain the fading curves of diffraction intensity and *N*_*e*_ values (doses that cause the diffraction intensity to drop by *1/e* or about 37% (44, 45)) of the samples at high cumulative electron doses under different imaging conditions. Comparative analysis shows that the fading curves and *N*_*e*_ values of C_44_H_90_ crystals do not change at different imaging electron dose rates, indicating that their radiation damage does not exhibit a dependence on the dose-rate effect. Under constant conditions, the fading curves and *N*_*e*_ values of C_44_H_90_ crystals are essentially the same in both conventional continuous and ultrafast pulsed modes. We find that, different from recent report (17), time-modulated pulsed electron beams do not seem to mitigate the electron radiation damage to the samples.

## Results

### The development of biological cryo-UEM and problem solving during experiment

A 200 kV biological cryo-UEM system with picosecond stroboscopic and nanosecond single-pulse electron mode was firstly developed, which was mainly composed of an ultrafast laser system and a biological cryo-EM system organically combined after modification (*SI Appendix*, Fig. S1). In the ultrafast laser system, two lasers were placed in the optical path, where the nanosecond laser was used in the single-pulse mode and the femtosecond laser was used in the stroboscopic mode. Considering the sensitivity of biological samples to laser absorption wavelengths, the femtosecond laser was also equipped with an optical parametric amplifier (OPA) to provide a wavelength tuning range of 630-2500 nm for the system. In the cryo-EM facility, the mature lens system, cryo-transfer system, cryo-box device, low-dose technology and real-time high-sensitivity data recording system of the commercial cryo-EM were used to obtain real-space and diffraction images for imaging in single-pulse mode and stroboscopic mode. The key points of such new cryo-UEM system were on the modification of the cryo-EM electron source and sample chamber, the precise introduction of the ultrafast laser, the co-axis adjustment in the pulsed mode, the search for the time-zero and ultrafast signals, the precise monitoring and adjustment of the laser position, and the installation and debugging of the direct electron detector. The cryo-UEM can be used to study the ultrafast dynamic conformational changes of photosensitive proteins. Here, it is mainly used to study the radiation damage behavior of soft matter represented by saturated hydrocarbon crystals (C_44_H_90_) under time-modulated pulsed electron beam, to explore its potential application in further mitigating the electron radiation damage of biological samples in cryo-EM imaging.

The C_44_H_90_ monocrystals laid flat on carbon membranes were prepared *via* the drop-casting method, and had a single-molecule-layer thickness with a rhombic morphology (Fig. 1). The diffraction patterns had a certain degree of defocusing to avoid detector saturation (46). A dark image for the correction of diffraction intensity was taken by masking the sample with shutter after each set of data was collected in both conventional continuous and ultrafast pulsed modes. The fading curves and *N*_*e*_ values of four diffraction points of {110} index in C_44_H_90_ monocrystals under electron radiation were statistically analyzed (Fig. 1). During the experiments, various problems such as the saturation of detector readings, the intensity drift of dark images, the instability of pulsed electron beam, and the temperature intolerance of samples in the cryo-UEM system, have brought difficulties to our experimental progress. The saturation of detector readings was mainly caused by the central transmission spot and diffraction spots being too “sharp” in diffraction mode, resulting in the intensity of these areas exceeding the detector reading range. We solved this problem by testing the defocus conditions required for the detector reading to be unsaturated before data collection (*SI Appendix*, Fig. S2). The intensity drift of dark images meant that the intensity of dark images taken by the detector, in the absence of electron radiation, was not stabilized around the zero value, but slowly drifted towards negative values over time. Since the dark image was taken to subtract background noise after the data collection of samples was completed (by default, the intensity of the dark image remained constant throughout the collection process), such drift effect would make the intensity value of the sample image obtained too high, ultimately resulting in an overestimated cumulative electron dose. We addressed this problem by performing multiple corrections on the dark image for its drift over time before data acquisition (*SI Appendix*, Fig. S3). The instability of pulsed electron beam involved two unfavorable aspects: first, data collection of fading curves in ultrafast pulsed mode usually took a long time, during which the intensity of the laser-excited pulsed electron beam would fade rapidly, losing the ability to diffraction imaging of the sample. The fading of pulsed electron beam was mainly caused by the adsorption of impurities on the cryo-UEM filament and the shift of probe laser in the filament position. We solved this problem by adding a new ion pump near the filament area and setting up a manual adjustment experimental strategy of “photoexcitation-thermal excitation-photoexcitation” (*SI Appendix*, Fig. S4-7). Second, the instability of pulsed electron beam also made it impossible to directly read the electron dose acting on each image. To solve such problem, we designed an experimental and calculation method that used Beer-Lambert’s law (44, 47) to obtain the average electron dose rate acting on the last image, and iterated forward in sequence to obtain the electron dose of all images. The feasibility and correctness of the data processing method were verified by experimental data in the conventional continuous electron-beam mode. During the experiment, we also noticed that after lowering the temperature of the samples in cryo-UEM chamber from 300 K to 100 K and leaving for a period of time, the samples at 100 K experienced the spontaneous fading of diffraction points, appearance of new amorphous rings and disappearance of morphological information (*SI Appendix*, Fig. S8), indicating the adsorption and condensation of impurities on the samples, which meant that the samples in cryo-UEM system could not tolerate a temperature of 100 K. By establishing a monitoring method for the state of samples at different temperatures in cryo-UEM, we finally confirmed that the lowest tolerable temperature of samples in the cryo-UEM chamber was ∼140 K (*SI Appendix*, Fig. S9). See the “Materials and Methods” section for more details.

**Fig. 1.**
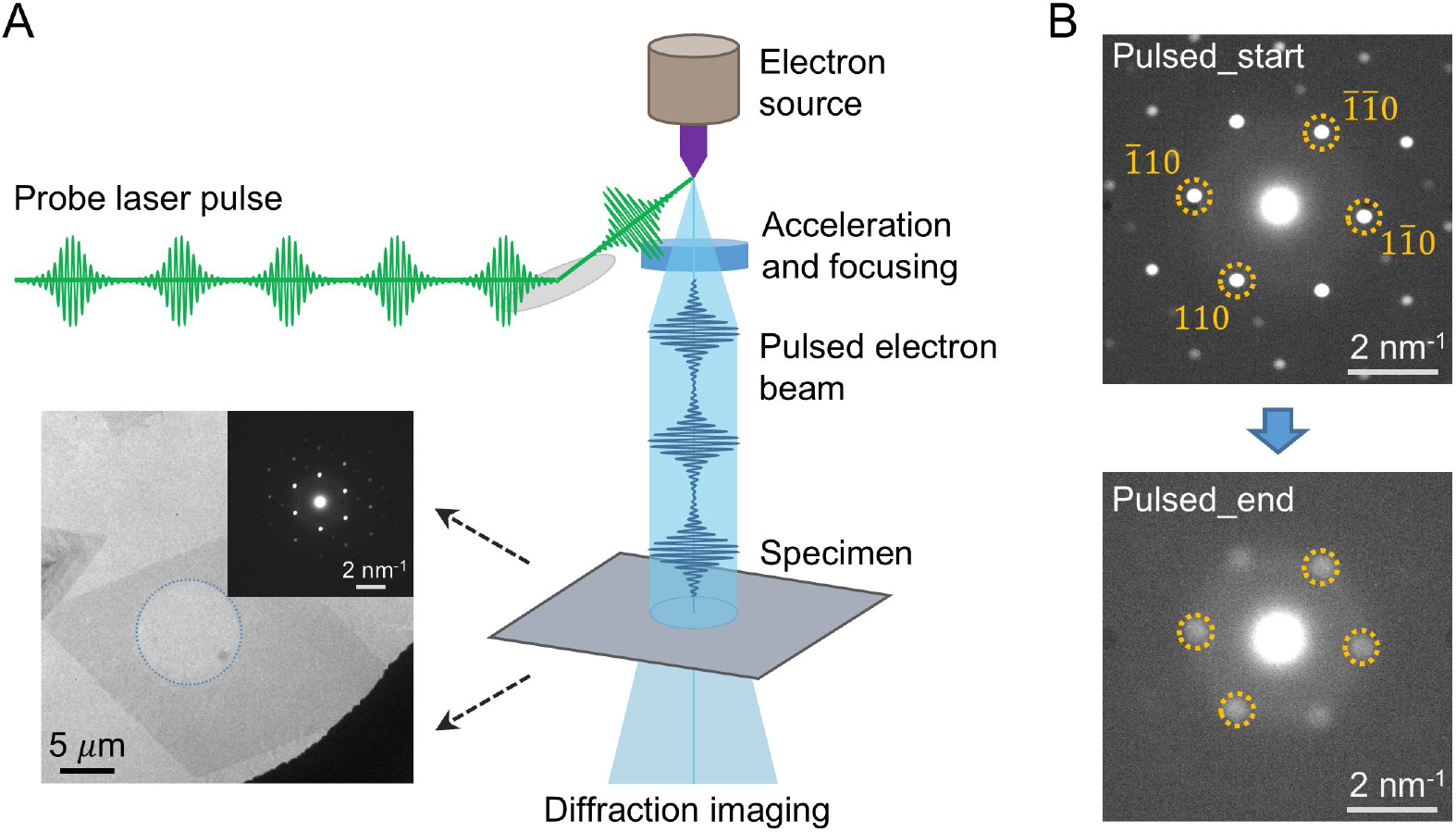
Experimental procedures for ultrafast pulsed electron imaging on C_44_H_90_ crystals. (*A*) Schematic diagram of pulsed electron imaging driven by an ultrafast laser system, with the radiated region on the rhombic crystal marked by a blue dashed line in the inset. (*B*) Diffraction patterns of C_44_H_90_ crystals obtained in the ultrafast pulsed mode, with the four diffraction points of {110} index used for the statistics of diffraction intensity marked by orange dashed lines.

### The radiation damage behavior of C_44_H_90_ crystals in conventional continuous electron-beam mode

The diffraction-intensity fading curves of C_44_H_90_ crystals were first measured at room temperature (300 K) and in conventional continuous electron-beam mode (200 kV) (Fig. 2A and *SI Appendix*, Fig. S10). The imaging electron dose rates were controlled by modulating the magnitude of cryo-UEM filament current and the spread of C2 condenser aperture. After each adjustment, a stabilization period was necessary to ensure the equipment reached a steady state, and data collection commenced upon confirmation of the stable imaging electron dose rates. The experimental results showed that, at different imaging electron dose rates in the range of 6.72×10^−5^ ∼ 1.00×10^−2^ e^−^/Å^2^/s, the fading curves of diffraction intensity of the C_44_H_90_ crystals were all essentially the same, exhibiting consistent decreasing trends and curve slopes. The *N*_*e*_ values obtained from the fading curves of diffraction intensity at different dose rates were also statistically analyzed, and as shown in Fig. 2B, the *N*_*e*_ values did not exhibit a correlation with the electron dose rates, which was a constant value of 3.62 ± 0.09 e^−^/Å^2^ in the conventional continuous electron-beam mode at 300 K. Furthermore, low-temperature experiments were also conducted. Despite the increased experimental complexity, the related experiments have following advantages: they enable both a lateral analysis of the diffraction-intensity fading curves and the *N*_*e*_ values of samples across different modes at the same temperature, as well as a longitudinal analysis of these parameters in the same mode at different temperatures. This can not only investigate the impact of temperature on the radiation damage behavior of samples under various modes, but also provide supporting evidence for the existence or non-existence of radiation-damage mitigation effect in the ultrafast pulsed mode. At different electron dose rates in the range of 8.23×10^−5^ ∼ 5.28×10^−3^ e^−^/Å^2^/s and 8.23×10^−5^ ∼ 1.89×10^−2^ e^−^/Å^2^/s, the fading curves of diffraction intensity of C_44_H_90_ crystals in the conventional continuous mode were measured at 200 K and 140 K, respectively (Fig. 2C-D). The experimental results showed that the fading curves of diffraction intensity of the C_44_H_90_ crystals at different electron dose rates were also essentially the same at 200 K, showing exactly the same decreasing trends and corresponding to a *N*_*e*_ value of 5.83 ± 0.12 e^−^/Å^2^. The fading curves of diffraction intensity, which were not correlated to the electron dose rates, were also observed at 140 K, corresponding to a *N*_*e*_ value of 9.75 ± 0.14 e^−^/Å^2^. These experimental results indicated that the radiation damage of C_44_H_90_ crystals under the electron beam exhibited no dependence on the imaging electron dose rates, but only a correlation with the cumulative electron doses.

**Fig. 2.**
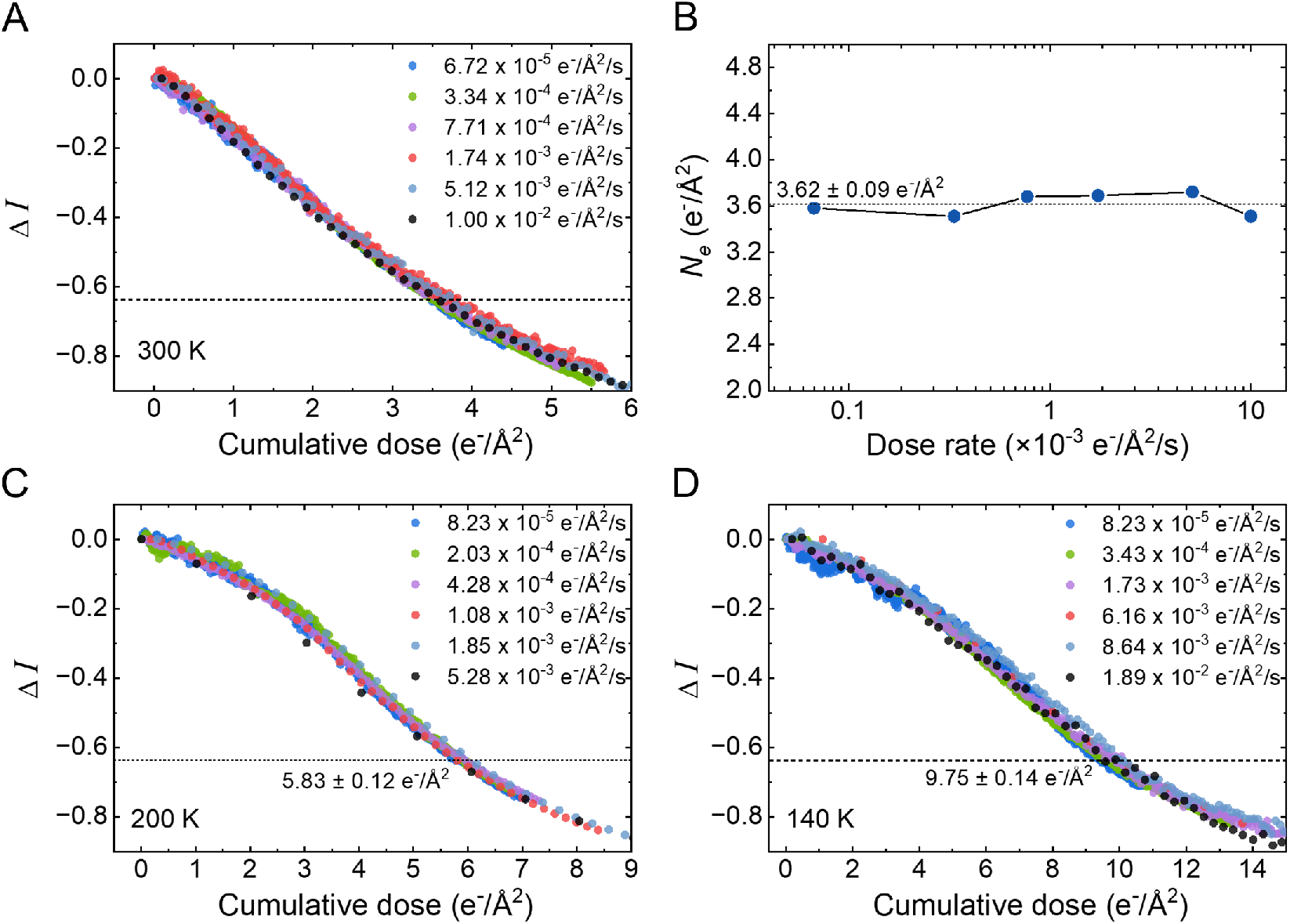
Electron radiation damage of C_44_H_90_ crystals in conventional continuous electron-beam mode and at different temperatures (300 K, 200 K and 140 K). (*A*) The fading curves of diffraction intensity on C_44_H_90_ crystals measured at different electron dose rates at 300 K, which showed the uncorrelation with electron dose rates. (*B*) The *N*_*e*_ values obtained at different electron dose rates in (*A*), respectively. The *N*_*e*_ values do not change with increasing electron dose rates and have a value of 3.62 ± 0.09 e^−^/Å^2^ within the systematic error. (*C-D*) The fading curves of diffraction intensity measured at different dose rates at 200 K (*C*) and 140 K (*D*). The corresponding *N*_*e*_ values are 5.83 ± 0.12 e^−^/Å^2^ and 9.75 ± 0.14 e^−^/Å^2^, respectively.

Moreover, compared to those at 300 K, the corresponding *N*_*e*_ values of C_44_H_90_ crystals at 200 K and 140 K were increased by ∼1.61 (5.83 *vs*. 3.62) and ∼2.69 times (9.75 *vs*. 3.62), respectively (Fig. 2), indicating the mitigation effect of radiation damage of the sample at low temperatures. In addition, we found that the fading curves of diffraction intensity at 200 K and 140 K showed a “first slow then fast” fading compared to those at 300 K. We attributed the “first slow then fast” two-stage form of fading curves to the “latent dose” effect (5, 48) exhibited by the samples at low temperatures: at low temperatures, the molecular fragments were “frozen” and their diffusion rates were suppressed. Only after reaching the latent dose, the stress generated in the crystals, due to the increasing number of fragments, would lead to the acceleration of the diffusion of the fragments (due to the cage effect (49), the molecular-diffusion rates were still lower than the rates at room temperature), resulting in a gradual fading of the diffraction intensity. We believed that the mitigation effect of radiation damage at low temperatures was produced synergistically by the presence of latent dose and the decrease in the diffusion rates of molecular fragments. Other experimental results related to the “latent dose” effect will be further analyzed in the following.

### The radiation damage behavior of C_44_H_90_ crystals in ultrafast pulsed electron-beam mode (200 kHz)

Using the newly established experimental strategy and data processing method (see the “Materials and Methods” section for details), the diffraction-intensity fading curves of C_44_H_90_ crystals in ultrafast pulsed electron-beam mode (laser repetition rate of 200 kHz) and in the temperature range of 140 ∼ 300 K were measured. The temporal modulation of the imaging electron beam by 200 kHz probe laser resulted in a 5 μs time interval between each two pulsed electron packets, to allow for the possible potential ground-state recovery and structural relaxation of samples. Fig. 3 showed the experimental results measured in the ultrafast multi-electrons-packet pulsed mode, where the number of electrons in each pulsed electron packet fluctuated between 1 and 8. More than three sets of data for each mode were measured independently and compared with the experimental results in the conventional continuous electron-beam mode (*SI Appendix*, Fig. S11). It can be found that the fading curves of diffraction intensity obtained at room temperature (300 K) were essentially the same for the multi-electrons-packet pulsed mode and the conventional continuous mode. At 200 K and 140 K, similar results to those in the conventional continuous mode were also observed in the ultrafast multi-electrons-packet pulsed mode. The *N*_*e*_ values of C_44_H_90_ crystals at different temperatures (300 K, 200 K and 140 K) and different imaging modes (conventional continuous and multi-electrons-packet pulsed modes) were further statistically analyzed. At 300 K, the *N*_*e*_ values of C_44_H_90_ crystals in the modes of conventional continuous and multi-electrons-packet pulsed were 3.62 e^−^/Å^2^ and 3.62 e^−^/Å^2^ respectively; at 200 K, the *N*_*e*_ values were 5.83 e^−^/Å^2^ and 5.76 e^−^/Å^2^ respectively; at 140 K, the *N*_*e*_ values were 9.75 e^−^/Å^2^ and 9.62 e^−^/Å^2^ respectively. These experimental results indicated that such multi-electrons-packet pulsed imaging mode could not mitigate the electron radiation damage of the C_44_H_90_ samples.

**Fig. 3.**
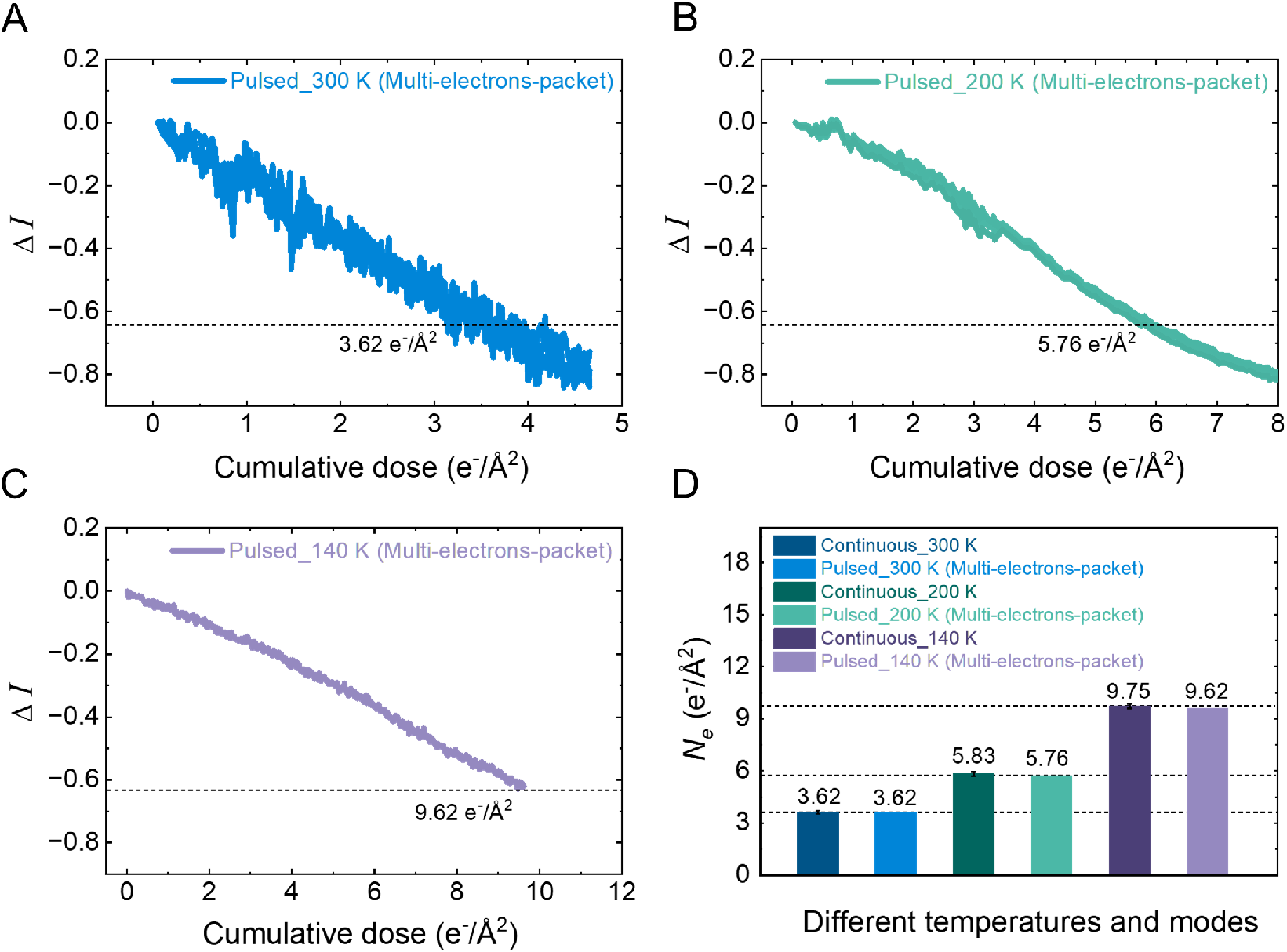
Electron radiation damage of C_44_H_90_ crystals in ultrafast multi-electrons-packet pulsed electron-beam mode at different temperatures. (*A-C*) The fading curves of diffraction intensity in ultrafast multi-electrons-packet pulsed mode at 300 K (*A*), 200 K (*B*) and 140 K (*C*), with *N*_*e*_ values of 3.62 e^−^/Å^2^, 5.76 e^−^/Å^2^ and 9.62 e^−^/Å^2^. (*D*) Statistics of *N*_*e*_ values of C_44_H_90_ crystals obtained in multi-electrons-packet pulsed mode and conventional continuous electron-beam mode at different temperatures.

In recent years, it has been reported that the mitigation effect of radiation damage in the ultrafast pulsed mode was related to the number of electrons per pulse (controlled by the laser pulse power) at the laser repetition rate of 200 kHz (17). Therefore, the diffraction-intensity fading curves of C_44_H_90_ crystals in near-single-electron-packet pulsed mode at the temperatures of 300 K, 200 K and 140 K, were further measured (in such mode, the number of electrons in each ultrafast electron pulse was about less than one (*SI Appendix*, Fig. S12)), and they were also compared with the fading curves obtained in the conventional continuous electron-beam mode. As shown in Fig. 4 and *SI Appendix*, Fig. S13, the experimental results showed that, whether at 300 K, 200 K or 140 K, the diffraction-intensity fading curves of the samples in such near-single-electron-packet pulsed mode were also essentially the same as those obtained in the conventional continuous mode, and had completely consistent curve forms. The *N*_*e*_ values of C_44_H_90_ crystals in the near-single-electron-packet pulsed mode and conventional continuous mode at different temperatures were also further statistically analyzed. At 300 K, the *N*_*e*_ values of C_44_H_90_ crystals in the modes of conventional continuous and near-single-electron-packet pulsed were 3.62 e^−^/Å^2^ and 3.67 e^−^/Å^2^ respectively; at 200 K, the *N*_*e*_ values were 5.83 e^−^/Å^2^ and 5.85 e^−^/Å^2^ respectively; at 140 K, the *N*_*e*_ values were 9.75 e^−^/Å^2^ and 10.04 e^−^/Å^2^ respectively. From these data, again no mitigation effect of radiation damage was observed to occur. In order to verify the experimental results, we repeated the test more than three times for each mode. These experimental results indicated that the ultrafast pulsed-electron imaging method with time modulation did not seem to mitigate the electron radiation damage of the samples, which was consistent with the experimental conclusion of “the independence of radiation damage and *N*_*e*_ values on imaging dose rates” obtained above.

**Fig. 4.**
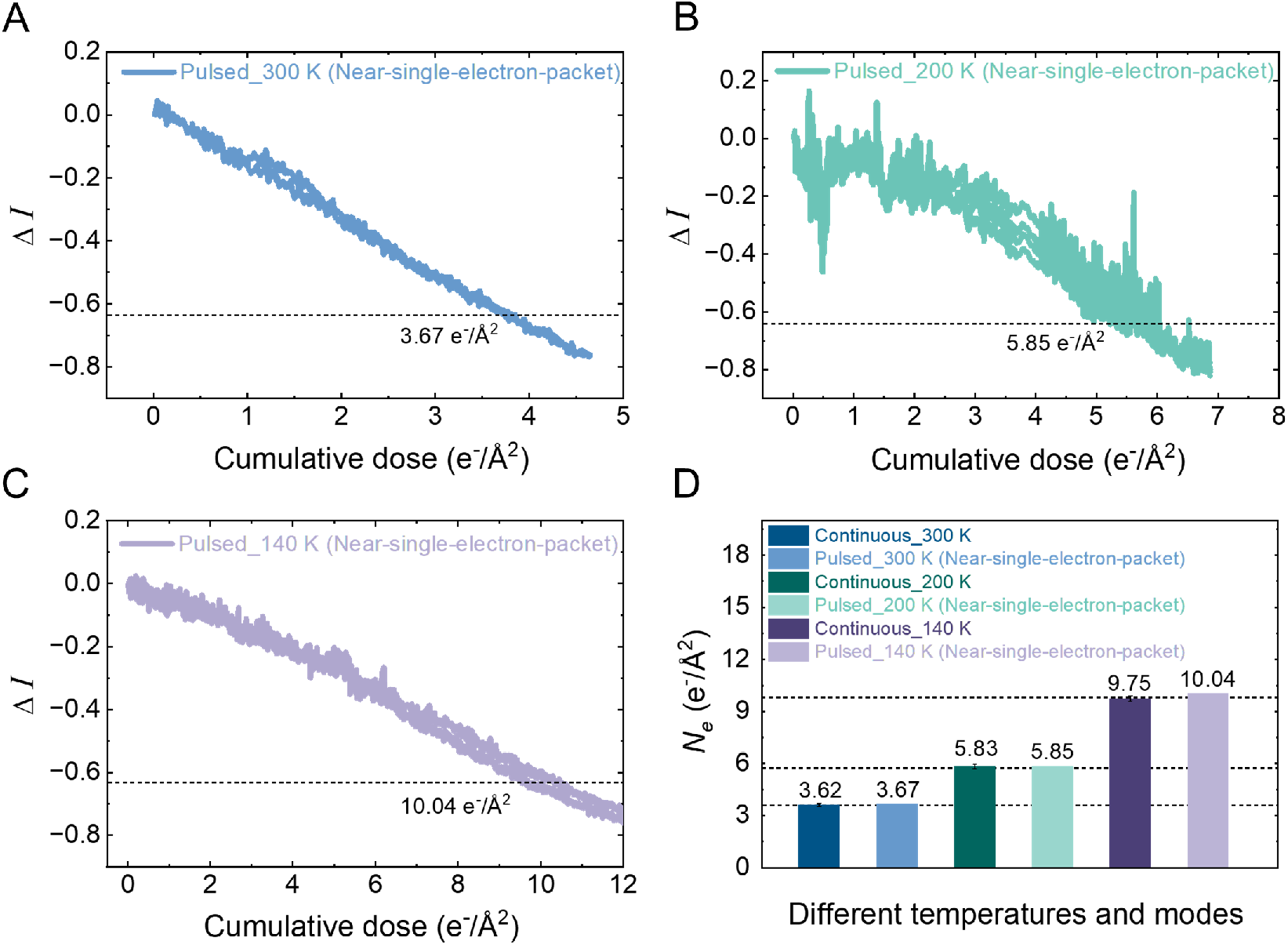
Electron radiation damage of C_44_H_90_ crystals in ultrafast near-single-electron-packet pulsed electron-beam mode at different temperatures. (*A-C*) The fading curves of diffraction intensity in ultrafast near-single-electron-packet pulsed mode at 300 K (*A*), 200 K (*B*) and 140 K (*C*), with *N*_*e*_ values of 3.67 e^−^/Å^2^, 5.85 e^−^/Å^2^ and 10.04 e^−^/Å^2^. (*D*) Statistics of *N*_*e*_ values of C_44_H_90_ crystals obtained in near-single-electron-packet pulsed mode and conventional continuous electron-beam mode at different temperatures.

### Experimental results at 100 kHz laser repetition rate and room temperature

To investigate whether the mitigation effect of radiation damage would be manifested at longer time intervals between pulsed-electron packets (controlled by the laser repetition rates), the fading curves of diffraction intensity of C_44_H_90_ crystals at 300 K and in pulsed mode were measured again by reducing the laser repetition rates from 200 kHz to 100 kHz. The 100 kHz laser repetition rate corresponds to a temporal interval of 10 μs between each electron pulse. Compared to the 200 kHz mode, the 100 kHz pulsed mode requires longer data collection time and lower pulsed electron dose rates (*SI Appendix*, Fig. S14). As shown in Fig. 5, at 300 K, the fading curves of diffraction intensity in multi-electrons-packet and near-single-electron-packet pulsed mode at 100 kHz were not different from the fading curves at 200 kHz, and the *N*_*e*_ values were 3.62 e^−^/Å^2^ and 3.58 e^−^/Å^2^, respectively. By statistical analysis of *N*_*e*_ values obtained in 100 kHz pulsed mode, 200 kHz pulsed mode and conventional continuous mode, it can be found that the *N*_*e*_ values under the three experimental conditions were basically the same (Fig. 5C). These data showed that modulating the number of electrons in the pulsed-electron packet and the time intervals between the pulsed electron packets did not affect the electron radiation damage of the samples. In other words, time-modulated ultrafast pulsed imaging did not appear to have a mitigation effect on radiation damage.

**Fig. 5.**
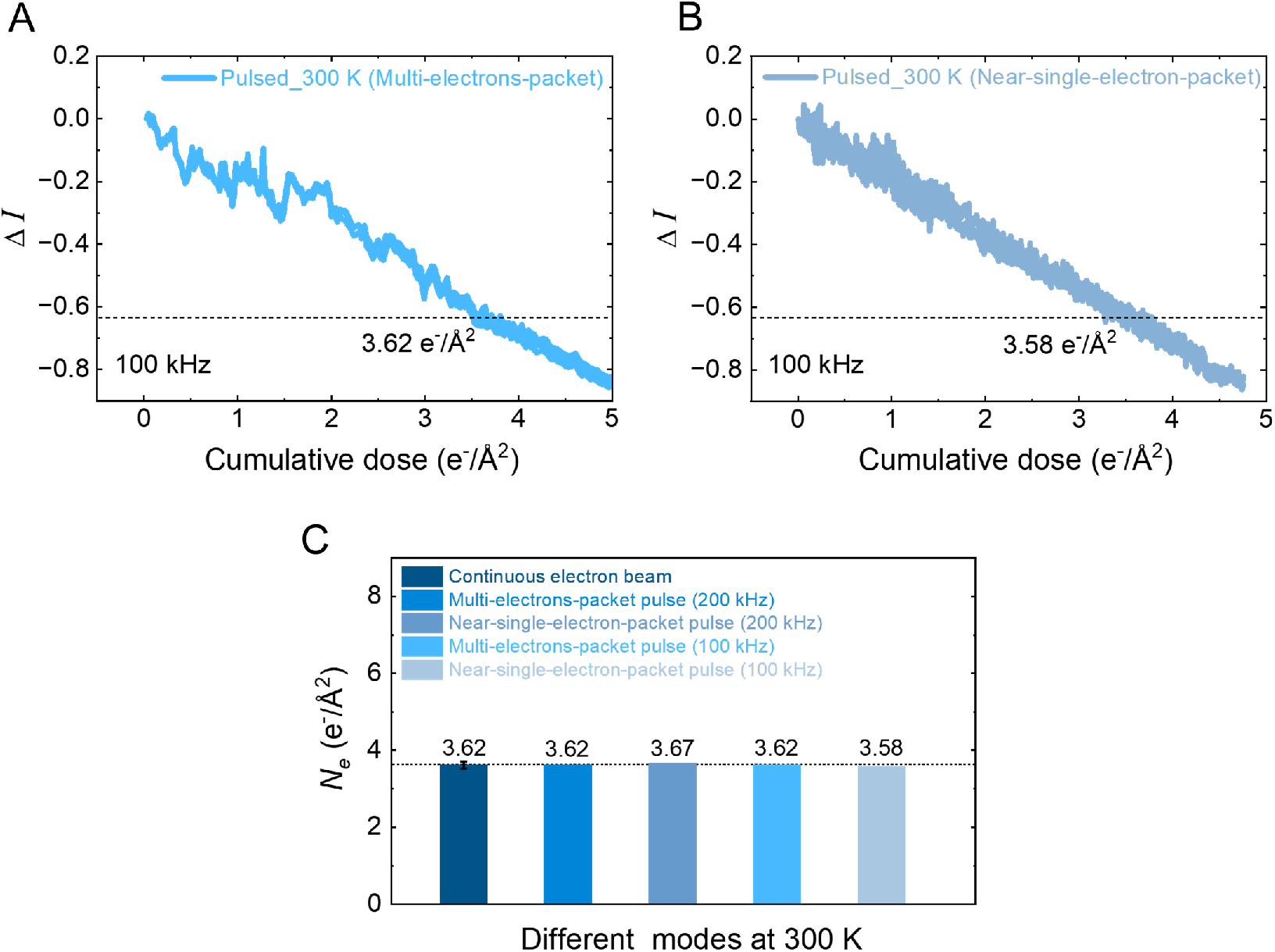
Electron radiation damage of C_44_H_90_ crystals at 100 kHz laser repetition rate and 300 K. (*A*) The fading curves of diffraction intensity in ultrafast multi-electrons-packet pulsed mode, with a *N*_*e*_ values of 3.62 e^−^/Å^2^. (*B*) The fading curves of diffraction intensity in ultrafast near-single-electron-packet pulsed mode, with a *N*_*e*_ value of 3.58 e^−^/Å^2^. (*C*) Statistics of *N*_*e*_ value of C_44_H_90_ crystals obtained under different imaging modes and laser repetition rates at 300K.

Moreover, a “latent dose” effect, similar to that reported by Siegel (48), was further observed on the C_44_H_90_ samples stored in a fume hood for a long time. At 200 K, the diffraction-intensity fading curves of the samples in both conventional continuous and ultrafast pulsed modes showed a greater degree of “first slow then fast” two-stage form, resulting in an increase in the *N*_*e*_ value of the samples compared with the original results (from 5.83 e^−^/Å^2^ to 8.26 e^−^/Å^2^) (*SI Appendix*, Fig. S15). Similar results were also observed at 140 K (the *N*_*e*_ values increased from 9.75 e^−^/Å^2^ to 13.90 e^−^/Å^2^) (*SI Appendix*, Fig. S16). An interesting finding was that compared with the original *N*_*e*_ values, the *N*_*e*_ values at 200 K and 400 K obtained on the current samples both increased by about 42% (5.83 *vs*. 8.26; 9.75 *vs*. 13.90). We noted that the mitigation effect of radiation damage was also not observed in the multi-electrons-packet and near-single-electron-packet pulsed modes on such samples.

## Discussion

The dose-rate effect and the mitigation effect of radiation damage in pulsed imaging mode have long been controversial. In recent years, the existence of mitigation effect of radiation damage in ultrafast pulsed mode for saturated hydrocarbon crystals has been reported at room temperature (17). However, perhaps due to the long data-collection time in pulsed mode and the challenges in maintaining the single-electron-packet pulsed mode, the author only measured the experimental data of pulsed mode with cumulative electron dose not higher than 0.1 e^−^/Å^2^ (only ∼2% of the *N*_*e*_ value). Here, through long-term and extensive data collection and repeated verification, we have carried out systematic and complete measurements of the diffraction-intensity fading curves of C_44_H_90_ crystals for the experimental conditions such as different imaging modes (conventional continuous and ultrafast pulsed), different temperatures (140 K to 300 K), different imaging dose rates (10^−5^ ∼ 10^−2^ e^−^/Å^2^/s) and different pulsed repetition rates (200 kHz and 100 kHz), and obtained the corresponding *N*_*e*_ values. The experimental results showed that the electron radiation damage of C_44_H_90_ crystals did not exhibit a correlation with the imaging electron dose rates, and no mitigation effect of radiation damage was observed in the ultrafast pulsed mode, whether at room temperature or low temperature.

We propose that the observed similarity in electron radiation damage caused by time-modulated ultrafast pulsed electron beams and continuous electron beams may be attributed to the following factors: (1) A single electron is sufficient to cause irreversible radiation damage to samples, such as bond breakage. In the recent experimental report, Konieczny *et al*. designed a chemical probe for free radical-mediated chain reactions, and found that a Dewar benzene crystal underwent up to 90,000 reactions per incident electron, greatly amplifying the ionization effect in diffraction mode (50). Moreover, Isaacson *et al*. systematically investigated the interaction between electrons and biomolecules under an electron microscope by characterizing the changes in electron energy loss spectra and electron diffraction patterns, and pointed out that the exponential decrease in the intensities of the energy-loss peak and diffraction peak was consistent with the “single hit” model (51), *i*.*e*., the electron radiation damage was produced by a single energy loss event rather than a succession of events (52); (2) The radiation damage of samples is only affected by the energy density, which has no time dimension. In many books on X-ray diffraction physics (53, 54), the following statements and related rigorous proofs can be found: the diffraction of crystals arises from scattered photons, and the scattering photons are exactly proportional to the energy density (photons/mm^2^), which has no time dimension. This means, the crystals are killed by energy density (cumulative dose), rather than time (55). Such statements are consistent with the experimental results that the dose-rate effect does not exist, *i*.*e*., the crystal quality obtained under the same cumulative dose is independent of the speed (electron dose rate) of the applied photons (electrons). On the other hand, as the oldest and most explanatory theory of the radiation-damage curves, the hit theory (51) mentioned in (1) first applied quantum physics ideas to biological problems. It is based on two physical observations and one postulate: I, ionizing radiation transmits the energy in the form of discrete packets. II, the interactions are independent of each other and follow a Poisson distribution. III, if a specified target has received a specified number of hits, the response under investigation will occur. From such statements, it is clear that again there is no consideration of the time dimension in hit theory; (3) Just as the dependence of the *N*_*e*_ values on the electron dose rates (dose-rate effect) was not observed in our experimental results, time-dependent cumulative effects, such as thermal effects, may contribute little to the electron radiation damage, implying that even if suitable pulse intervals are provided to allow for structural relaxation and recovery of the sample, the effect on the sample radiation damage is negligible. In a previous study, Thornburg *et al*. observed the warming of samples under a 100 kV electron beam by means of thin film thermocouples, and found that the temperature rise of the samples under “typical” operating conditions was less than 10 K, which was much lower than the empirical values previously reported. Even under extreme beam conditions (100 μA), the temperature of the samples increased by only about 15 K (56); (4) If cumulative effects, such as thermal effects, predominate in the radiation damage of saturated hydrocarbon samples, the mitigating effect of radiation damage may be manifested in a pulsed electron beam with suitable time modulation. However, the relaxation and recovery of radiation damage caused by thermal effects could not be accomplished within the time interval of 200 kHz or 100 kHz pulses (based on our experimental results). In addition, as Glaeser said (25), to a first approximation, the average time between electrons could be arranged to be the same whether the electrons were emitted from a continuous source or a pulsed source. Simply reducing the intensity of the standard continuous source, should provide a long enough pause between inelastic scattering events to allow for the assumed thermal relaxation and recovery. It means that if there is a radiation-damage mitigation effect in the pulsed electron-beam mode, then the mitigation effect should also be observed in the continuous mode with low electron intensity, but this is not the case; (5) The radiation-damage mechanism of samples under electron beam is complex and may be affected by the synergistic effect of multiple damage effects. While relevant experimental and theoretical reports have been emerging in recent decades, as Stenn and Bahr lamented in their early reviews (57), we have to admit that we still know very little about the radiation-damage mechanism of samples due to the extremely complex cascade and synergistic processes involved. Perhaps aside from the longitudinal cascade responses that occur over time, there are also many lateral and cross-propagating cascade responses of radiation damage to the sample at the same time. Given the difficulty of precisely monitoring the events occurring during radiation damage of samples, it still remains uncertain whether the intermittent cessation of electron beam on the microsecond time scale could lead to the mitigation or disappearance of damage effects such as chemical bond breaking or free radical generation.

It is noteworthy that, with the comprehensive advancement of relevant software and hardware in recent times, the issue of electron radiation damage to samples has emerged as a significant technical bottleneck hindering the further enhancement of material-conformation characterization techniques across various scientific fields, and we hope that more and more groups will join in the systematic study of the electron radiation damage behavior of samples. We believe that it is meaningful to reconsider this topic now, as it concerns the future development of electron imaging technology in new application environments and a new round of “resolution revolution”. Here, our research has opened up a new corner for the study of sample damage behavior under electron-beam radiation and the integration and proposal of related physical mechanisms. These findings underscore the intricate nature of radiation damage processes and highlight the need for further exploration of the fundamental principles governing damage mitigation under different experimental conditions.

In the next research work, using the newly established cryo-UEM equipment, we will integrate relevant technical methods to establish and improve the technical framework and experimental platform of TEM imaging for femtosecond-microsecond kinetic observation of biomolecules. With such established technique, potential intermediate transition states during photoexcitation of photosensitive proteins, will be investigated to provide detailed structural descriptions of the ultrafast photochemical events they undergo, and to elucidate the conformational pathways and the corresponding molecular mechanisms of the photosensitive proteins during photoexcitation (58). Compared with X-ray serial crystallography, such cryo-UEM technology has significant advantages of low equipment cost, high popularity, small amount of sample required, high data collection efficiency, mature data processing technology, and the potential for *in-situ* real-space imaging of amorphous samples, which is expected to revolutionize the entire research field.

## Conclusion

In conclusion, we built a cryo-UEM device based on the ultrafast laser system, and explored the radiation damage behavior of conventional continuous electron beam and ultrafast pulsed electron beam on C_44_H_90_ crystals. The experimental results show that the fading curves and *N*_*e*_ values of samples are not related to the imaging electron dose rates and do not show any dependence on the dose-rate effect. As the temperature decreases (300 to 140 K), the *N*_*e*_ values of the samples increase by 2.69 times (3.62 to 9.75 e^−^/Å^2^), indicating the cryoprotective effect on radiation damage to the samples. At the constant temperature, the fading curves of diffraction intensity and *N*_*e*_ values of the samples in both multi-electrons-packet and near-single-electron-packet pulsed modes are roughly the same as those in conventional continuous mode, even when the results are obtained at different pulsed repetition rates. Our experimental results suggest that the time-modulated pulsed electron beam do not seem to mitigate the electron radiation damage occurred on the samples, which is consistent with the views of Glaeser *et al*.(25). Such results indicate that ultrafast pulsed electron imaging with time modulation does not appear to be advantageous in developing into an effective means to overcome the radiation-damage problem of samples in cryo-EM technology. Our findings offer new insights and experimental basis for the radiation damage behavior of samples under electron beams, and provide guidance and inspiration for elucidating the fundamental principles of radiation damage.

## Acknowledgments

We thank Yujia Zhai for helpful discussions, Zhongwen Li and Chaoxiang Chen for modifying the cryo-UEM. The study is funded by the National Key Research and Development Program (2019YFA0904101), National Natural Science Foundation of China (31925026, T2495223, T2495223), the Strategic Priority Research Program of the Chinese Academy of Sciences (XDB37040102), National Key Research and Development Program (2021YFA1301500), Beijing Natural Science Foundation (JQ24056).

